# Comparative study on the effects of different types of acute exercise on cortisol levels and cognitive functions

**DOI:** 10.1101/2024.09.14.613054

**Authors:** Zhongshu shao, Talifu Zikereya, Jiazheng Peng, Minggang Luo, Jinliang Chen, Kaixuan Shi

**Affiliations:** Department of Physical Education, China University of Geosciences, Beijing, China; Department of Physical Education, University of Science and Technology Beijing

**Keywords:** acute exercise, cortisol, cognitive functions, orienteering

## Abstract

This study examines the different effects of dual-task and single-task exercises on cortisol levels and cognitive performance. Seventeen male college students participated, performing each exercise on separate days. Salivary samples were taken before and after exercise to assess cortisol levels. Cognitive functions were measured using the 2-Back Test, Bells Test, and Mental Rotation Test. Results showed that both exercises significantly improved reaction times across cognitive tests. However, dual-task exercise caused a more notable increase in cortisol levels (*p*=0.002) than the single-task. A negative correlation between post-exercise cortisol levels and reaction times suggests that higher cortisol levels might enhance cognitive processing speed. Cognitive performance ac-curacy remained unchanged across both exercises. These findings indicate that while both exercise types enhance cognitive speed, dual-task exercise triggers a greater cortisol response, potentially offering more benefits for cognitive processing speed without affecting accuracy. This research highlights the complex interaction between acute exercise, stress hormones, and cognitive function, providing insights into how different exercise types influence cognitive performance through hormonal pathways.

## 1. Introduction

Acute exercise is a common physiological stressor that can trigger the body’s stress response and have complex effects on both physiology and psychology[1]. The stress response induced by exercise is typically accompanied by significant physiological changes such as increased heart rate and blood pressure. Additionally, exercise can directly affect brain function by increasing cerebral blood flow and neurotransmitter release[2].

When the body is stimulated by acute exercise, the hypothalamic-pituitary-adrenal (HPA) axis is activated[3]. Cortisol, the end product of the HPA axis, plays a crucial role in maintaining homeostasis and coping with stress[4]. Changes in cortisol levels can be measured through blood or saliva samples and are therefore commonly used as biomarkers for assessing the stress response[5].

In addition to reflecting stress levels, cortisol is closely related to cognitive function. Appropriate levels of cortisol are thought to promote cognitive performance, such as improving memory and attention[6, 7]. However, excessively high or low cortisol levels may adversely affect cognitive function, such as causing memory decline and attention deficits[8, 9]. This suggests that there may be an inverted U-shaped relationship between cortisol and cognitive function, with moderate levels of cortisol being most beneficial for cognitive performance[10].

Different types of exercise can have bidirectional regulatory effects on cognitive function through cortisol. Angelidis found that in acute cognitive task performance, the anxiety significantly increased cortisol levels, especially under the condition of high cognitive load[11]. Additionally, Zschucke discovered that during single-task exercise, participants in the exercise group showed a significant reduction in cortisol response to the Montreal Imaging Stress Task (MIST). This reduction may be due to acute exercise mitigating cortisol response to stress through a negative feedback mechanism in the hippocampus and prefrontal cortex[12]. Furthermore, Davis found that compared to single-task exercise, dual-task exercise might lead to greater competition for cognitive resources, as it involves simultaneous physical exercise and cognitive tasks, potentially causing a short-term decline in cognitive performance[13]. However, there is still a lack of in-depth research on the effects of single-task and dual-task exercise on cortisol levels and cognitive function[14]. Although existing research has found that exercise, stress, cortisol, and neural activity are interrelated and can influence cognitive function to some extent, it remains unclear to what extent cortisol affects cognitive function under different types of exercise and whether these different types of exercise lead to distinct cortisol response patterns[15, 16]. The conflicting results further complicate the cognitive and motor effects. However, the specific advantages of activities like orienteering, which require both physical exertion and cognitive skills, it is an ideal model to study movement in dual-task mode may offer new perspectives for addressing this issue. Here, we observed changes in cognitive function in relation to cortisol levels and examined whether cortisol modulates cognitive function to enhance performance after an exercise intervention. We utilized single-task (5000m running) and dual-task (orienteering) exercises, involving two different sports, to assess their effects on cortisol levels and cognitive performance. Cortisol levels were measured before and after the exercise to assess their impact on cognitive function.

## 2. Materials and Methods

### 2.1 Participants

This study included seventeen male college students from a university in Beijing, who were non-physical education majors. All participants were orienteering enthusiasts with over one year of training experience, aged 18 to 22 years. The participants’ average training duration was 7-12 h/week; their average BMI was 20.76 ± 2.39, and all participants had a BMI within the normal range.; all were right-handed and had no history of any diseases. The criteria for selecting subjects included age between 19.00 ± 3.00 years; body mass index (BMI) of 22.00 ± 5.00 kg/m^2^; no smoking or alcohol consumption, and no use of various drugs; good physical condition, and no interference from various neurological or chronic diseases (such as insomnia or various cardiovascular diseases, metabolic diseases, etc.) in this study. The recruitment for this study began on December 3, 2024, and the experiment concluded on February 5, 2024. All participants have signed the paper version of the informed consent form. This research plan was approved by the Ethics Committee of China University of Geosciences.

### 2.2 Experimental design

The study employed a cross-sectional design with within-subject controls conducted over three days, comprising three independent experimental stages (Figure 1).

**Figure 1.**
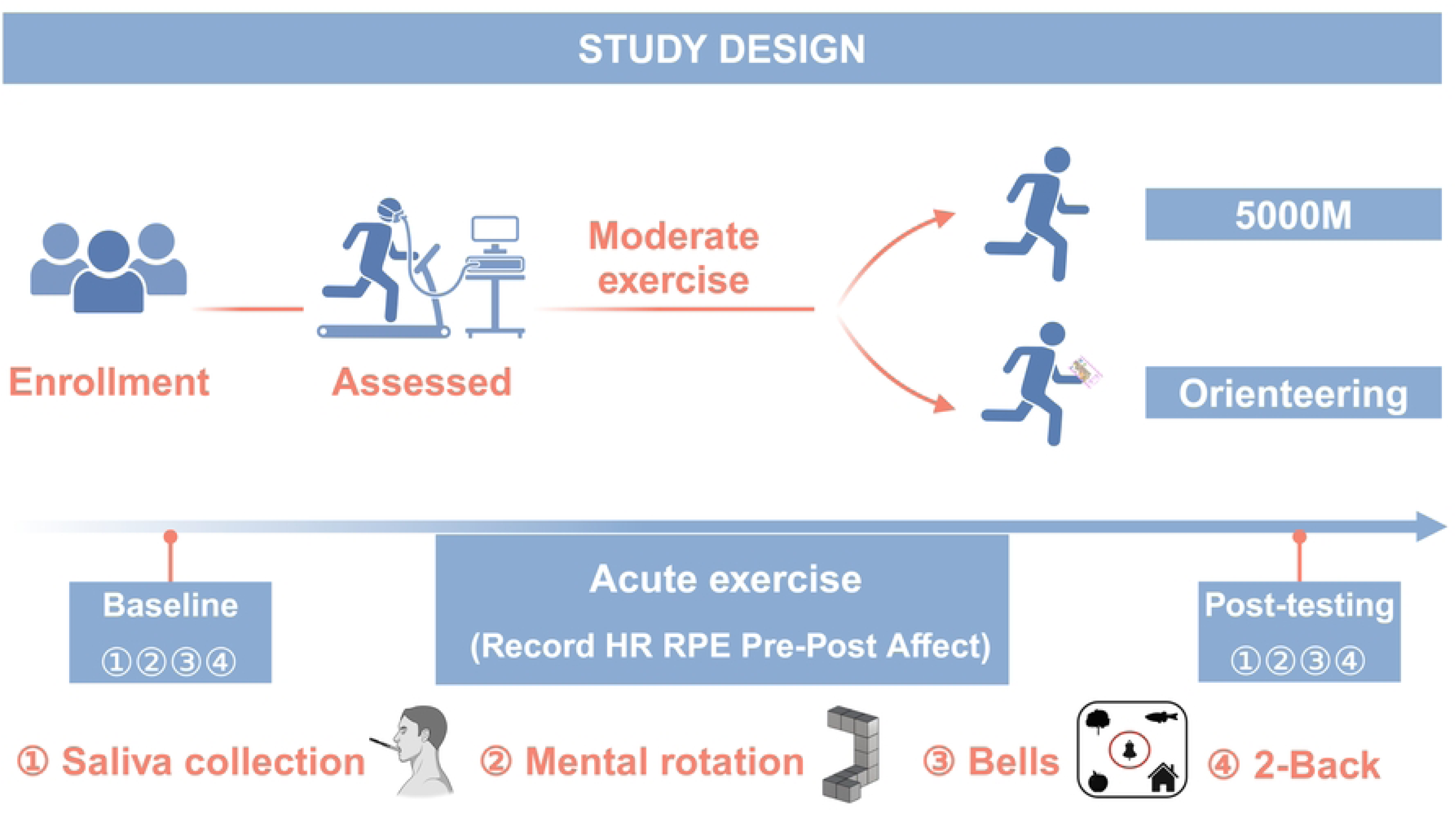
Experimental Design. This figure illustrates the overall experimental design used in the study. Participants underwent cognitive testing, body composition measurement, maximal oxygen uptake, and salivary cortisol collection. This was followed by 5000m and orienteering tests on different days, followed by salivary cortisol collection and cognitive function testing immediately after completion.

The first session: Participants arrived at the laboratory in the morning, adjusted the chair height and angle, rested for 10 minutes, and then completed cognitive function tests. After completion of the test, Body Composition was tested using the InBody 570 Body Composition Analyzer (InBody Co., Ltd., Seoul, Korea). Finally, the maximal oxygen uptake test was performed.

The second sessions: Participants arrived at an unfamiliar venue and performed a short-distance orienteering test. The orienteering map, designed with input from domestic experts, with the total course length being around 5000 meters. It was a short-distance orienteering practice, not a race, and required them to complete the exercise at moderate intensity. The terrain was a park with slopes, flat roads, woods, and ponds, and the entire training process could last about 40-50 minutes, including a warm-up time of about 8-10 minutes. Participants wore heart rate monitors (Polar H10) and GPS devices to track their heart rates, running routes, and distances during the test. Participants used a compass and map to navigate the orienteering course. Upon completing the exercise, their heart rates and running routes were recorded, and salivary cortisol levels were measured immediately. Then, cognitive function tests were immediately performed[17].

The third sessions: Seven days after the second day of testing, participants were required to complete a 5000m aerobic running exercise on a standard track. Similarly, they were also asked to complete the exercise at moderate intensity, participants needed to wear heart rate monitors to record their heart rate (Polar). After completion, the participants’ heart rates were immediately recorded, and salivary cortisol was collected. Immediately after the collection, cognitive tests were performed. Each time after completing the orienteering and 5000m running, participants underwent the Borg Rating of Perceived Exertion (RPE) scale test to determine the exercise intensity.

To control the influence of the human circadian rhythm, all experiments were conducted between 9:00 and 11:00 in the morning, and the subjects avoided consuming caffeine, alcohol, etc., 12 hours before the experiment and avoided eating food 60 minutes before the experiment.

### 2.3 Maximal oxygen uptake test

Following body composition assessment, participants completed a maximal oxygen uptake test according to the Bruce protocol. Participants used a Cosmed K5 Gas Analyzer (Cosmed Srl, Rome, Italy) during treadmill running, with the workload increasing every 3 minutes until maximum intensity could no longer be sustained despite encouragement. During the test, heart rate (HR) was monitored and recorded using a Polar heart rate monitor to measure peak heart rate. Peak work capacity (pVO2 peak) was measured, and VO_2_max was estimated using the highest 20-second average achieved before exhaustion to ensure participants’ abilities were at the same level[18]. Running and orienteering exercise intensity was determined by %HRmax. In the current study, LI (Low intensity) exercise was defined as 40% VO_2_max (∼63% HRmax), MI (Moderate intensity) as 60% VO_2_max (∼75% HRmax), and HI (High intensity) as 85% VO_2_max (∼91% HRmax)[19]. We used the RPE scale in combination with heart rate to determine the exercise intensity rating.

### 2.4 Cognitive Function Test

Cognitive performance was evaluated using three sport-specific tests: working memory (2-Back Test), visual attention (Bells Test), and mental rotation ability (Mental Rotation Test). These three abilities were tested using the 2-Back Test, The Bells Test, and the Mental Rotation Test, respectively. The 2-Back task and Mental Rotation task were completed on a computer, with participants seated approximately 1m from the screen and the chair height adjustable for comfort. The Bells Test was completed using paper and pencil.

2-Back test: Participants were required to memorize the location of sequentially displayed stimuli. After each presentation, they had to determine whether the current stimulus matched the position of the stimulus shown two steps earlier and respond accordingly by pressing the correct or incorrect key.

Bells test: A sheet of paper with 35 target figures (bells) and various distractors (horses, fish, trees, apples, keys) was used. Each row contained 5 bells and 40 distractors. Participants had to circle as many bells as possible with a pencil within 30 seconds. At the end of the experiment, the participant’s score was the number of correctly circled bells. The Bells test was chosen because it can assess participants’ visual search ability[20].

Mental Rotation test: The Mental Rotation Test, commonly used in neuroscience, assesses participants’ spatial visualization and reasoning abilities. Participants were asked to observe 48 pairs of 3D objects, mentally rotate them, quickly determine whether the object could be reduced to a marker by rotation, and respond as quickly as possible by pressing a key representing correct or incorrect[21]. Orienteering requires athletes to determine their location and direction of travel in unfamiliar environments using maps and compasses. During this process, athletes must adopt a self-centered perspective, viewing themselves as an object within the map’s coordinate system and continuously adjusting their position and orientation to reach the destination. Therefore, mental rotation ability is an essential skill for orienteering athletes[22].

### 2.5 Cortisol collection and analysis

Enzyme-linked immunosorbent assay (XY-9H0641, Shanghai, China) was used to detect salivary cortisol levels, with the kit purchased from Shanghai Xin Yu Bioengineering Research Institute. Saliva samples were collected in tubes, centrifuged within 2 hours to separate saliva, and the supernatant was analyzed using an enzyme-linked immunoassay detector. Throughout the process, participants were not allowed to consume stimulants (such as coffee, chocolate, etc.) within three hours before the test. They were required to abstain from drinking and eating for 60 minutes and rinse their mouth with clean water. Before collection, participants were checked for staying up late, oral ulcers, oral wounds, or dental caries that could affect sample accuracy.

### 2.6 Statistical analysis

SPSS26.0 was used for statistical analysis. All results were expressed as mean ± standard deviation (SD). Pearson was used to verify the correlation of the data. p ≤ 0.05 was considered statistically significant. The differences of cortisol variables between the two groups before and after exercise were analyzed by independent sample t test. The data of cognitive function in resting state, after orienteering and after 5000m exercise were analyzed. Repeated measures ANOVA was used to compare the changes of cortisol in the three states between the two groups. Figure by GraphPad Prism9.5 drawing.

## 3. Results

### 3.1. Participants

Table 1 summarizes the main characteristics of the participants. Participants’ characteristics included an average age of 19.41 ± 1.03 years, weight of 62.76 ± 7.68 kg, height of 173.82 ± 3.85 cm, body fat percentage of 12.53 ± 4.33%, BMI of 20.76 ± 2.39 kg/m², and VO_2_MAX of 58.91 ± 6.31 ml/kg/min. Designed to control and monitor the exercise intensity of the subjects. After the exercise by filling the RPE scale and records the heart rate, determine the movement of moderate intensity.

**Table 1.** Basic characteristics of the participants (Mean ± SD; n = 17)

| Variables |  | Mean (SD) |
| --- | --- | --- |
| Age (year) | | $19.41 \pm 1.03$ |
| Weight (kg) | | $62.76 \pm 7.68$ |
| Height (m) | | $173.82 \pm 3.85$ |
| Body fat (%) | | $12.53 \pm 4.33$ |
| BMI (kg/m<sup>2</sup>) | | $20.76 \pm 2.39$ |
| VO<sub>2</sub>MAX (ml/kg/min) | | $58.91 \pm 6.31$ |
| HR (bpm) | Dual-task | $157.88 \pm 19.11$ |
| | Single-task | $158.00 \pm 14.47$ |
| RPE | Dual-task | $14.67 \pm 0.83$ |
|  | Single-task | 13.88 ± 0.76 |

<sup>1</sup> This table summarizes the main characteristics of the participants, including their age, weight, height, body fat percentage, BMI, VO<sub>2</sub>MAX, heart rate during dual-task and single-task exercises, and RPE for both activities. BMI = Body Mass Index; HR = Heart Rate; RPE = Borg Rating of Perceived Exertion.

### 3.2. Cortisol

Figure 2A and Table 2 present the results of paired t-tests conducted to evaluate cognitive function indices in participants at rest, and following both dual-task and single-task exercises. Results revealed a significant elevation in cortisol levels after dual-task exercise, with the paired data showing a robust correlation (*p* = 0.002; t = 3.83). In contrast, no significant change in cortisol levels was observed after single-task exercise (*p* = 0.054; t = 2.07).

**Figure 2.**
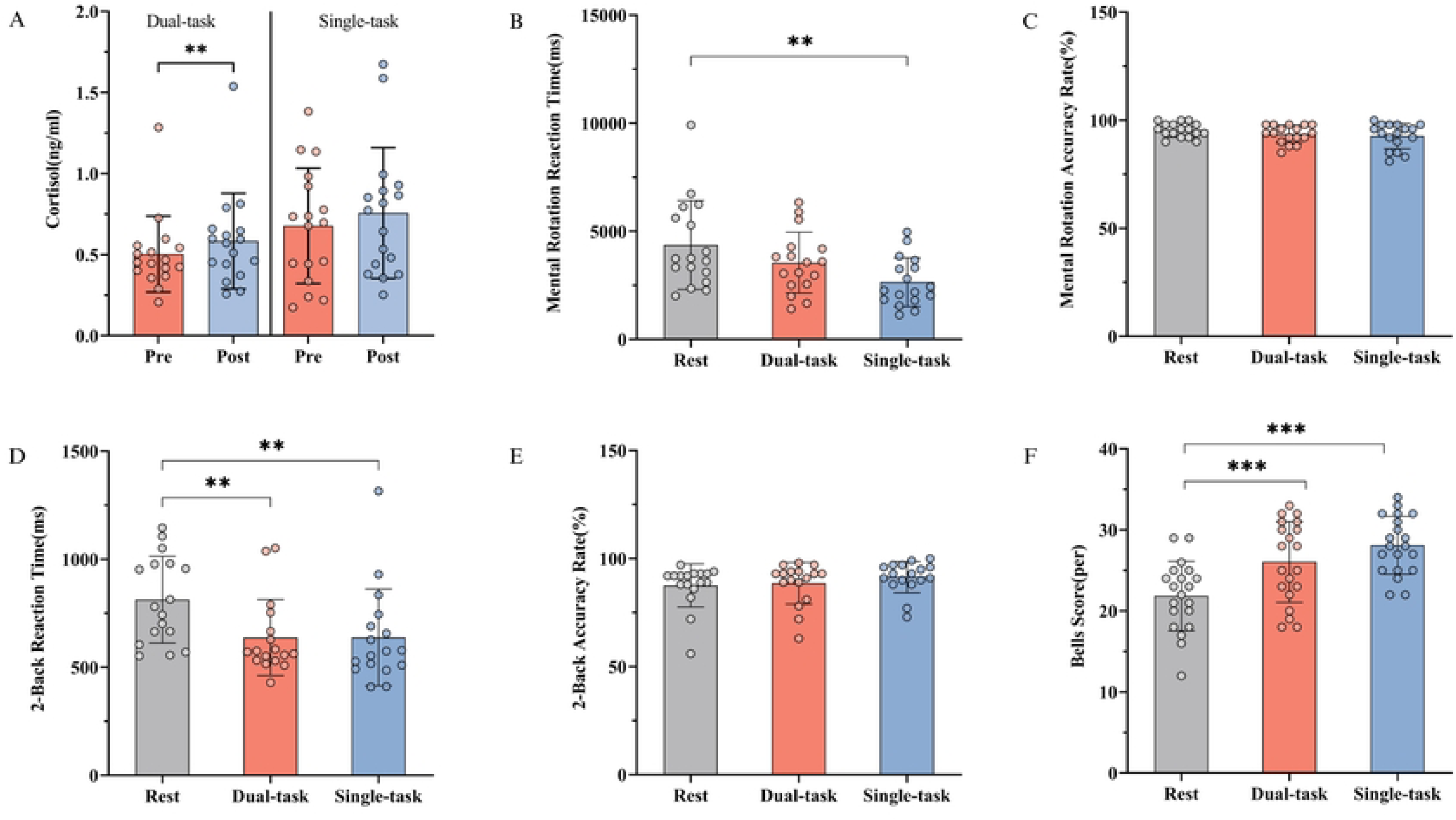
Cortisol levels and cognitive function test results. A: Changes in salivary cortisol levels in subjects before and after dual-task and single-task exercises. B-C: Mental rotation reaction time and accuracy in different states. D-E: 2-Back reaction time and ac-curacy in different states. F: Bells Score in different states. *: *P* < 0.05; **: *P* < 0.01; *** *P* < 0.001.

**Table 2.**
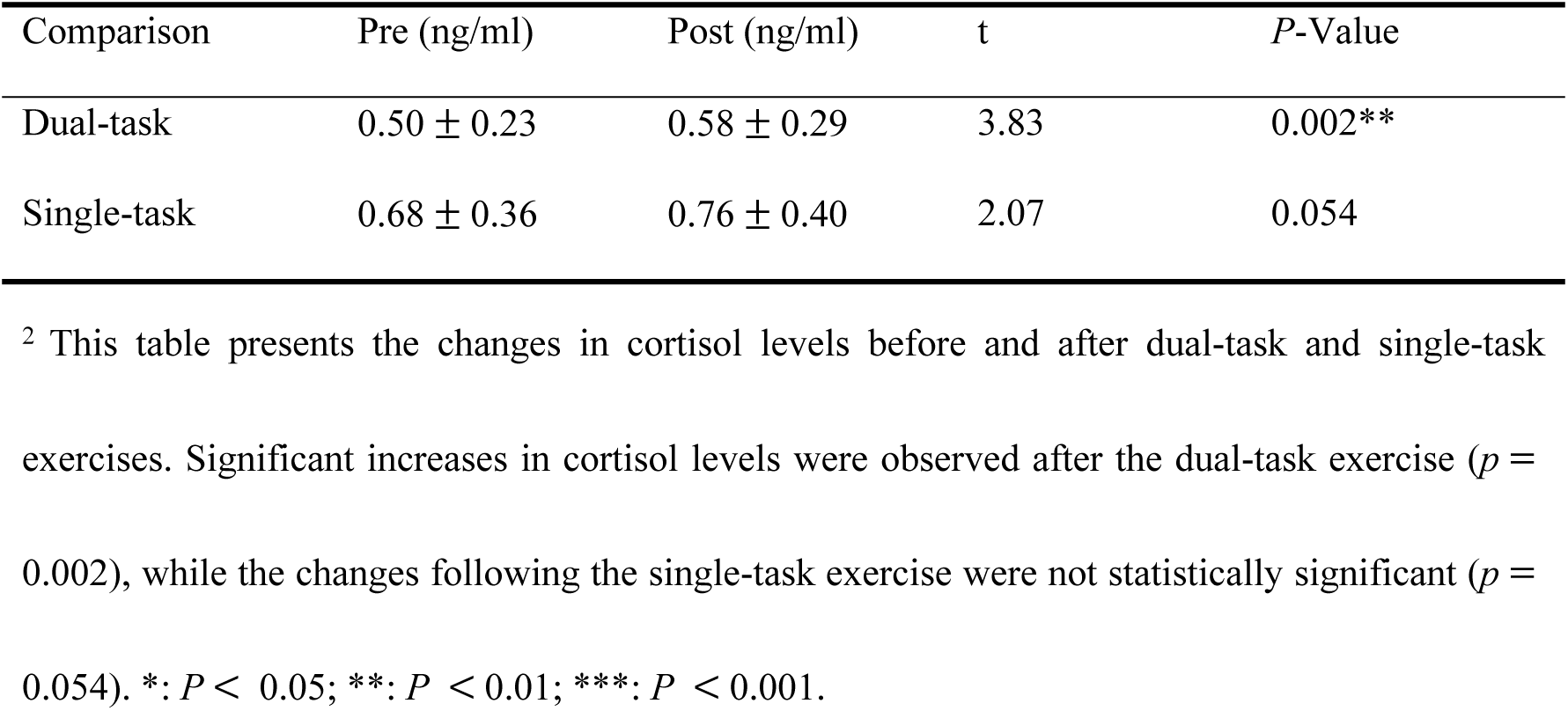
Changes in cortisol levels before and after exercise.

### 3.3. Cognitive Function

#### 3.3.1. Mental Rotation

A repeated measures analysis was used to assess cortisol and mental rotation test results across different conditions (Figure 2B; C and table 3). It was found that there was a significant change after single-task exercise compared with that at rest (F = 5.73 *p* = 0.008; R^2^ = 0.26).

**Table 3.**
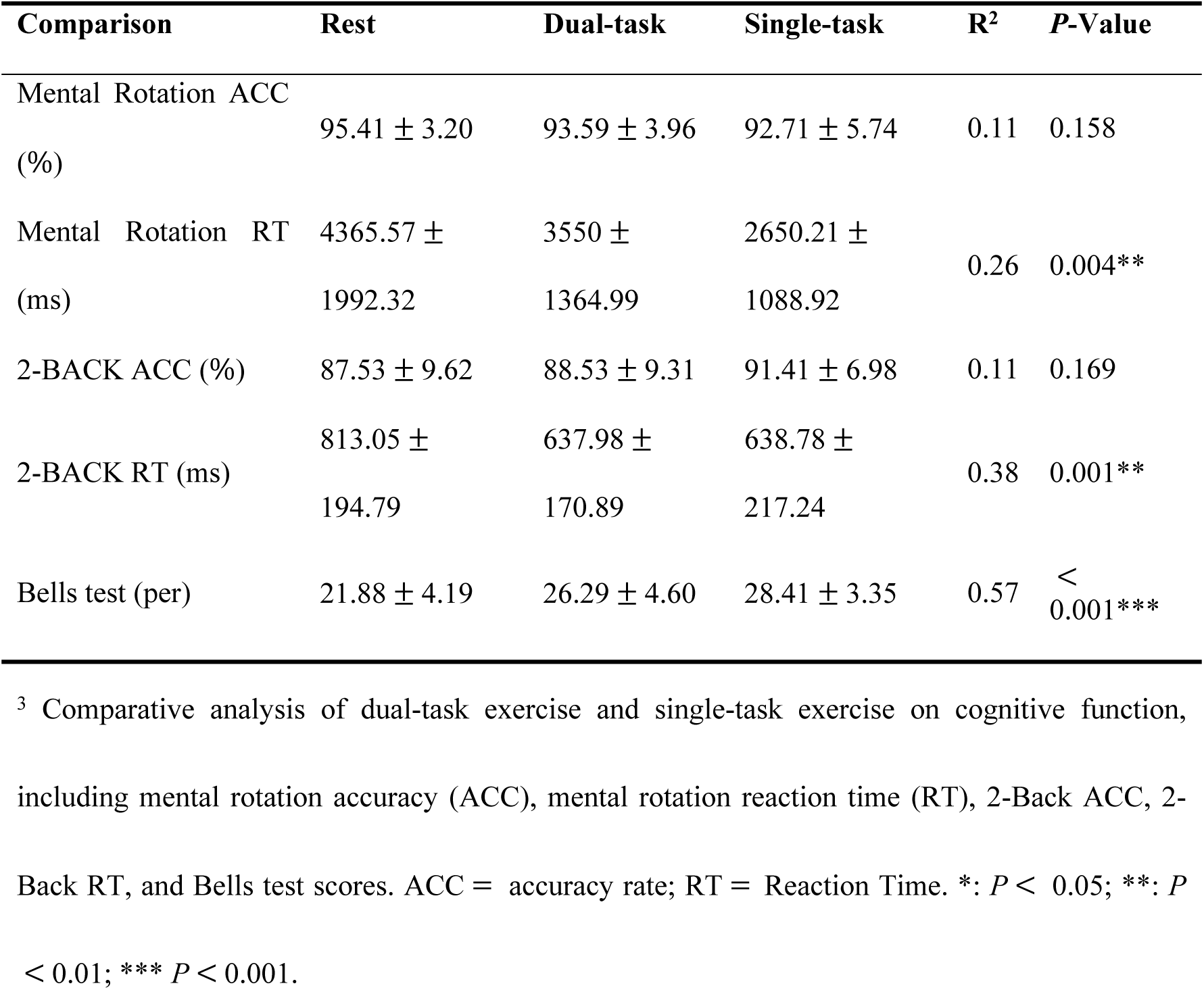
Comparative Analysis of Acute Exercise Types on Cognitive Function Responses.

Multiple comparisons showed that although reaction times decreased after dual-task exercise compared to the resting state, this difference was not significant (*p* = 0.221; t = 1.61). In contrast, single-task exercise significantly reduced reaction times compared to the resting state (*p* = 0.004; t = 3.38).

One-way ANOVA revealed no significant differences in accuracy between the groups, with R² accounting for 10.90% of the variance (F = 1.96; *p* = 0.158; R² = 0.11). Multiple comparisons indicated that neither dual-task (*p* = 0.360; t = 1.31) nor single-task exercise (*p* = 0.119; t = 1.94) led to significant changes in accuracy compared to the resting state.

#### 3.3.2. 2-Back Test

Data from Figure 2D; E and table 3 were analyzed using a one-way ANOVA on reaction times and accuracy in the 2-Back Test across three distinct conditions. The analysis showed a statistically significant difference among the groups. Post-hoc tests compared the two types of exercise to the resting state. Reaction time significantly decreased after dual-task exercise compared to the resting state (*p* = 0.001, t = 3.81). Similarly, reaction time significantly decreased after single-task exercise compared to the resting state (*p* = 0.001, t = 3.80). For accuracy, the one-way ANOVA showed no significant differences between the groups (F = 1.88, *p* = 0.169, R^2^ = 0.11). Post-hoc tests indicated no significant changes in accuracy for either dual-task exercise (*p* = 0.866, t = 0.48) or single-task exercise (*p* = 0.137, t = 1.87) compared to the resting state.

#### 3.3.3. Bells Test

The Bells test data for the three different conditions were compared using a one-way ANOVA (Figure 2F and table 3). The score analysis revealed a statistically significant difference between the two groups (F = 24.89, *p* < 0.001; R^2^ = 0.57).

Multiple comparison tests were conducted to compare the two different types of exercise with the resting state. The scores significantly increased after dual-task exercise compared to the resting state *(p* < 0.001; t = 4.65). Similarly, the scores significantly increased after the single-task exercise compared to the resting state *(p* < 0.001; t = 6.92).

### 3.4. Correlation between Cortisol and Cognitive Function

The relationship between cortisol levels and cognitive functions was analyzed using Pearson’s correlation coefficient following dual-task and single-task exercises (Table 4, Figure 3). Results showed that after dual-task exercise, cortisol levels were significantly negatively correlated with reaction times in the Mental Rotation Test (r = -0.45, *p* = 0.008) but not with accuracy (r = 0.23, *p* = 0.191). The Bells Test scores showed a positive but non-significant correlation with cortisol (r = 0.30, *p* = 0.087). In the 2-Back Test, reaction time was significantly negatively correlated with cortisol levels (r = -0.43, *p* = 0.012), while accuracy did not show a significant correlation (r = 0.17, *p* = 0.350).

**Figure 3.**
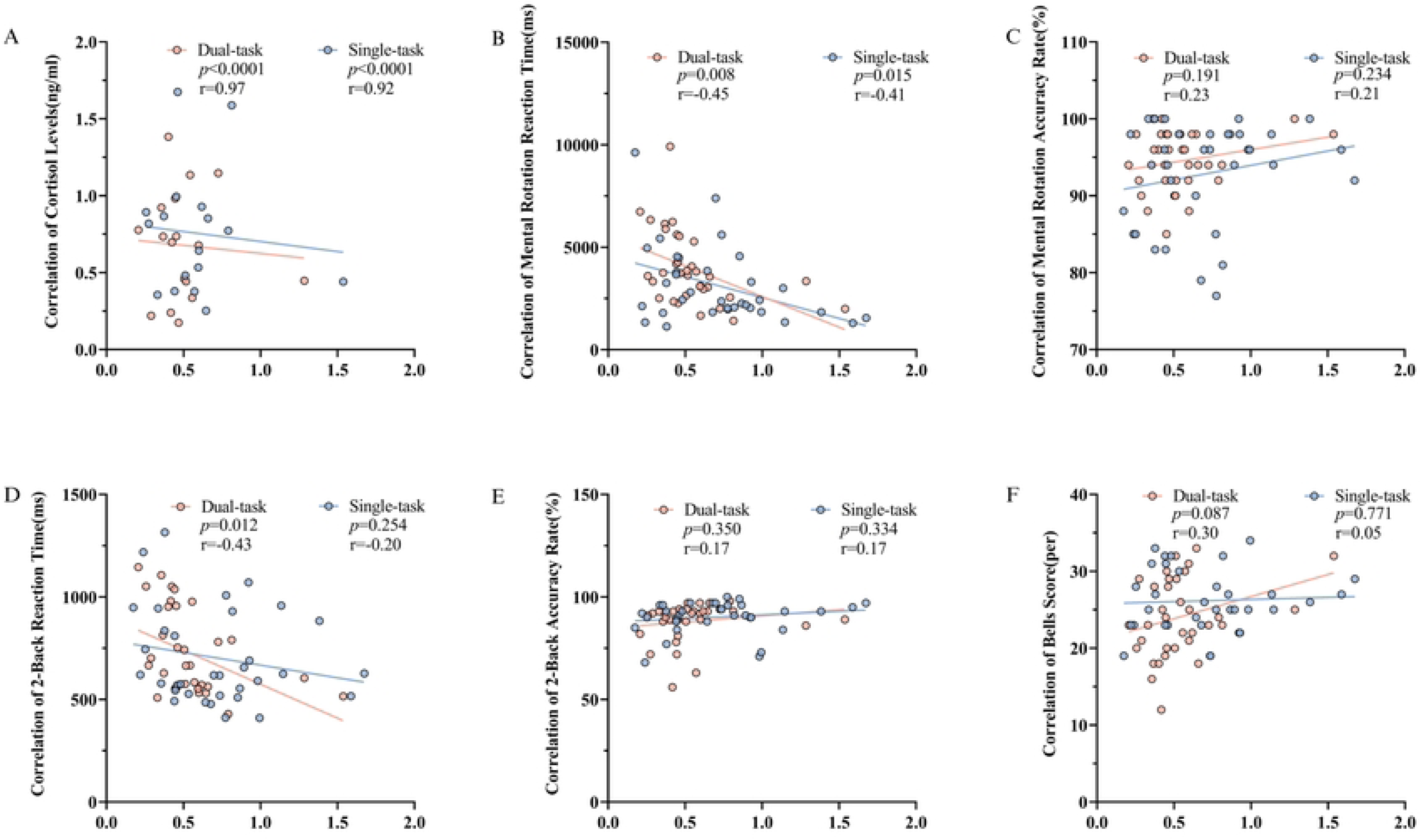
Correlation between Cortisol Levels and Cognitive Function Post-Exercise. A: Correlation of Cortisol Levels (ng/ml). B-C: Correlation of Mental Rotation Reaction Time (ms) and Accuracy Rate (%). D-E: Correlation of 2-Back Reaction Time (ms) and Accuracy Rate (%). F: Cor-relation of Bells Score (per).

**Table 4.** Correlation between different types of exercise, cortisol and cognitive function.

| Comparison | Variables | *P*-Value | *r* |
| --- | --- | --- | --- |
| Cortisol of Orienteering | Mental Rotation ACC | 0.191 | 0.23 |
|  | Mental Rotation RT | 0.008** | -0.45 |
|  | 2-BACK ACC | 0.350 | 0.17 |
|  | 2-BACK RT | 0.012* | -0.43 |
|  | Bells score | 0.087 | 0.30 |
| Cortisol of 5000m | Mental Rotation ACC | 0.234 | 0.21 |
|  | Mental Rotation RT | 0.015** | -0.41 |
|  | 2-BACK ACC | 0.334 | 0.17 |
|  | 2-BACK RT | 0.254 | -0.20 |
|  | Bells score | 0.771 | 0.05 |
| Cortisol of Orienteering pre | Cortisol of Orienteering post | < 0.001*** | 0.97 |
| Cortisol of 5000m pre | Cortisol of 5000m post | < 0.001*** | 0.92 |

<sup>4</sup> This table shows the correlation between cortisol levels and cognitive function indices (mental rotation accuracy, mental rotation reaction time, 2-Back accuracy, 2-Back reaction time, and Bells test scores) before and after orienteering and a 5000m run. Significant negative correlations were observed between cortisol levels and reaction times for both exercises ( $p < 0.01$ ), indicating that higher cortisol levels were associated with faster reaction times. \*: $P < 0.05$ ; \*\*: $P < 0.01$ ; \*\*\* $P < 0.001$ .

Following the single-task exercise, the reaction times in the Mental Rotation Test again demonstrated a significant negative correlation with cortisol levels (r = -0.41, *p* = 0.012), while accuracy correlations remained non-significant (r = 0.21, *p* = 0.23). The Bells Test results showed a very weak and non-significant correlation with cortisol (r = 0.05, *p* = 0.770). Both reaction time and accuracy in the 2-Back Test exhibited non-significant correlations (r = -0.20, *p* = 0.254 for reaction time; r = 0.17, *p* = 0.334 for accuracy).

Overall, our findings suggest that cortisol levels may be associated with certain cognitive function test scores, particularly in reaction times, following physical activities.

## 4. Discussion

This study aims to explore the effect of dual-task and single-task exercise on cortisol levels and cognitive functions under acute exercise. The results show that dual-task exercise significantly increased cortisol levels compared to single-task exercise. Both types of exercise improved reaction times in cognitive tasks without significantly affecting accuracy. These findings contribute to understanding how different types of exercise regulate stress responses and cognitive function.

Cortisol is a measure of psychological stress biomarkers and is therefore called a “stress hormone.” It is released in response to physical or social psychological pressure sources[23, 24]. Acute exercise stimulates an increase in cortisol, resulting in neurochemical changes[15]. Pastor et al. found that acute exercise can improve student cognitive function[25]. In contrast, prolonged exercise training is frequently associated with decreased cortisol levels, likely due to stress adaptation mechanisms elicited by sustained repetitive physical activity[26, 27]. Additionally, prolonged exercise may reduce basal cortisol levels and the response to stress by modulating the sensitivity of the HPA axis, enhancing homeostasis, and improving the body’s ability to adapt to stress, thus reducing the negative effects of chronic stress on cognitive function[12].

Our findings confirmed that dual-task exercise under acute exercise conditions significantly increased cortisol levels, whereas single-task exercise did not significantly alter cortisol levels. This discrepancy may result from the stronger stress response activated by the complex cognitive load in dual-task exercise. Dual-task exercise, which requires simultaneous physical and cognitive activity, places a greater demand on the activation of the HPA axis, resulting in higher cortisol release[2].

In addition, our findings also unveil the acute movement under the condition of the intricate relationship between cortisol and cognitive function (Figure 4). Despite significantly elevating cortisol levels, dual-task exercise concurrently enhances reaction time during cognitive tasks without compromising accuracy[28, 29]. Certain studies have indicated that acute increases in cortisol enhance attention and speed of information processing, thereby augmenting short-term cognitive performance. This may be attributed to the positive impact of cortisol on alertness and cognitive processing in response to acute stressors[30, 31]. The activation of the hypothalamic-pituitary-adrenal (HPA) axis during exercise plausibly explains these findings. Acute exercise acts as a stressor, prompting the hypothalamus to release corticotropin-releasing hormone (CRH). This, in turn, stimulates the pituitary gland to secrete adrenocorticotropic hormone (ACTH), which triggers the adrenal cortex to produce cortisol[12, 32, 33]. Increased cortisol levels enhance alertness and cognitive function, potentially explaining the improved reaction times observed in our study. In dual-task exercise, the combined physical and cognitive demands may further elevate the HPA axis response, resulting in higher cortisol levels compared to single-task exercise[3, 33]. Nevertheless, further investigations are warranted to ascertain whether this transient cognitive enhancement can be sustained over a prolonged duration[34–36].

**Figure 4.**
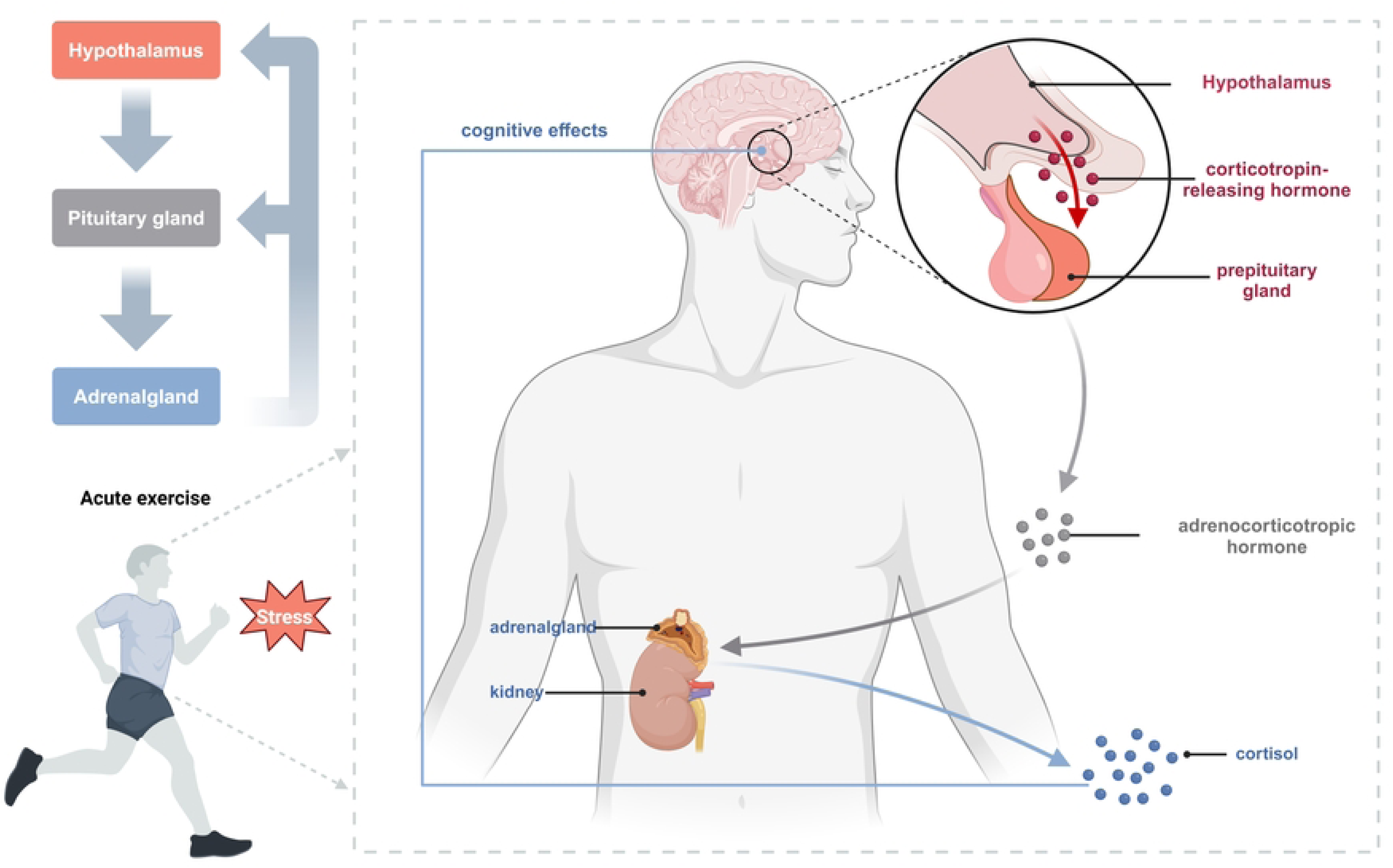
Mechanism of Cortisol Response to Acute Exercise. This figure showed that the mechanism by which acute exercise influences cortisol secretion through the hypothalamic-pituitary-adrenal (HPA) axis. Stress induced by exercise triggers the hypothalamus to release corticotropin-releasing hormone (CRH), which activates the pituitary gland to secrete adrenocorticotropic hormone (ACTH). ACTH then stimulates the adrenal glands to release cortisol. The increase in cortisol has cognitive effects, such as the regulation of stress responses[41–43].

The effect of dual-task exercise on cognitive function remains a contentious issue. On one hand, dual-task exercise simultaneously engages physical and cognitive abilities, leading to competition for cognitive resources. This competition can transiently impair cognitive function, especially when executing complex tasks or under fatigue[37–39]. Conversely, other studies have shown that cognitive reaction times decrease and working memory capacity improves following dual-task exercise. This improvement may be related to the participants’ exercise duration and intensity, as well as specific cognitive demands[13, 40]. Our findings align with the latter observation. This outcome may be due to our participants being university students with prior orienteering experience, which provides them with stronger cardiopulmonary fitness and specialized cognitive advantages related to orienteering. Additionally, both activities were conducted at moderate intensity. Thus, while cortisol levels increased during dual-task exercise, accuracy was not compromised, and reaction times improved.

Although this study provides new insights into the effects of acute exercise on cortisol and cognitive function under single- and dual-task conditions, it has some limitations. First, the sample size is small and comprises only male university students, which may limit the generalizability of the results. Future studies should consider larger and more diverse samples to further validate these findings. Second, this study examined only two specific types of exercise. Future research should explore the effects of other types of exercise on different cognitive functions. Additionally, incorporating data from long-term exercise interventions would help to more comprehensively understand the regulatory effects of exercise on cortisol and cognitive function.

## 5. Conclusions

This study indicates that dual-task acute exercise provokes a higher cortisol response compared to single-task exercise. Both forms of exercise significantly improved reaction times in cognitive tasks without significantly affecting accuracy. This suggests that moderate-intensity acute exercise, whether single- or dual-task, can enhance cognitive processing speed and alertness. Future research should expand the participant population and exercise types to further investigate the relationships between exercise, stress responses, and cognitive function.

## Author Contributions

Conceptualization, Zhongshu shao, Talifu Zikereya and Kaixuan Shi; Data curation, Jiazheng Peng; Methodology, Zhongshu shao, Talifu Zikereya and Kaixuan Shi; Project administration, Minggang Luo and Jinliang Chen; Validation, Jiazheng Peng; Writing – original draft, Zhongshu shao and Talifu Zikereya; Writing – review & editing, Kaixuan Shi.

## Institutional Review Board Statement

The study design complies with the Declaration of Helsinkiethical standards. The protocol was reviewed and approved by the Ethics Committee of China University of Geosciences (CUBG-EC-2023002).

## Informed Consent Statement

All participants signed a written statement of informed consent.

## Acknowledgments

In addition to the listed authors, assistance was provided by Yichen Yang and Qiangwei Wang.

